# Geneplot: a coordinate conversion approach for graphical representation of protein domain data on the exon-intron structure of a gene

**DOI:** 10.1101/2022.11.08.513416

**Authors:** Daniel Gonzalez-Ibeas

## Abstract

Graphical representation of single gene data, including subgenic features, polymorphisms and protein domains, is part of the regular routine of genome analyses. In the case of protein-coding genes, integration of such information with the exon-intron structure has advantages since intron polymorphisms may also have a biological impact, and the extent to which exons and protein domains overlap is of interest to evolutionary research. This report introduces *geneplot*, an open-source Python library to generate this type of graphical output from standard file formats. The library applies a coordinate conversion approach in order to represent protein domain data on genomic areas.

## Introduction

Since the advent of next generation sequencing technologies, hundred of reference genomes from different types of organisms have been made publicly available. After the assembly of the draft sequence, annotation is the next key step to identify structural elements with biological function. Nowadays, emerging interest is on the rise about non-coding RNA genes and transposons due to the impact they may have on the genome and cellular dynamics, but protein-coding genes have been traditionally the main target of annotation projects. Protein domains represent structural units that confer particular functionalities and very often self-fold during protein maturation. Protein domain information obtained from popular repositories and tools [1] is crucial for the functional annotation of the gene and to understand the protein product. In bioinformatic charting, these domains are typically represented on the protein sequence, but less often on the full gene sequence. However, the subgenic exon-intron structure provides valuable information in order to study alternative splicing (which mediates functional diversification of conserved genes), exon orthology relationships (consequence of gene evolution during speciation events), and intron persistence on genes among different lineages [2], [3]. Thus, integration of domain information on the exon-intron topology becomes a relevant approach to study the interplay between gene structure and function. For example, correlation between domain topology and exons among several genes of eukaryotes has been an active area of research that has fed the exon shuffling theory, as a mechanism to explain how domains could be reused during evolution [3]–[5].

Additionally, single nucleotide polymorphisms (SNPs) are one of the preferred molecular markers, because they are abundant, highly polymorphic and genome-wide distributed in nearly every organism [6]–[9]. Many SNPs within the coding area of genes have been reported to contribute to phenotypic diversity, including human diseases or agricultural traits. SNPs overlapping with protein domains have been always considered stronger candidates to identify trait-associated polymorphisms, even promoting the development of dedicated databases [10], [11]. However, introns may contain functional elements whose mutations can influence the biological fitness of an organism, as exemplified in the human genome [12], highlighting the relevance of including exon-intron topology in the graphical representation of SNP data.

Several visualizing tools exist to display genetic information at the single gene level. Most of them allow a straightforward representation of SNPs and other data types, but they don’t often integrate protein domain characteristics on the exon-intron structure, and finding a tool to undertake this task becomes challenging, especially in non-model organisms. For example, online genome browsers, such as the UCSC (https://genome.ucsc.edu/index.html, [13]), are valuable general-purpose resources that provide protein domain information mapped on genic subfeatures, but they are limited to the model species listed in its database. In the UCSC database, Pfam domains are firstly identified on protein sequences and after they are mapped to the transcripts themselves using a specialized tool called *pslmap*. As an alternative to this mapping approach, programming languages bring also different options to convert the coordinates of transcript/protein sequences to genome positions (and vice versa) from raw data sources, such as genome annotation files or protein annotation software. Then, converted coordinates can be used in your favorite visualizing tool. Some R packages allow this avenue, for example *ensembldb* and *GenomicFeatures* [14], [15]. In principle, they are intended to be used only with Ensembl, UCSC and BioMart databases, but a customized database from a GFF or GTF file can be created by the *GenomicFeatures* library, allowing the possibility to be applied in non-models species. Likewise, Perl also offers a solution to this problem with the *Bio::Location* library of *Bioperl* [16]. However, at the time of writing, there is only a *Mapper* module under development in Python to get this goal up, but not fully integrated in the latest stable release of *Biopython* [17]. This publication introduces *geneplot*, a Python library for integrating and visualizing exon-intron topology, SNP data and protein domain information. It implements its own coordinate conversion method within a Python framework and it can be applied to non-model species, since it takes as input standard files and formats generated from popular tools, including InterproScan [18], the General Feature Format v3 (GFF3, https://github.com/The-Sequence-Ontology/Specifications/blob/master/gff3.md) and the Variant Call Format (VCF, https://github.com/samtools/hts-specs).

## Implementation

The library combines information from genome annotation files in GFF3 format, the standard output of InterproScan containing domains identified on protein sequences, and SNP data from VCF files (Figure 1). The first step is the instantiation of a ‘genome’ object as a Python class using local paths to the files of annotation, Interpro domains and polymorphisms as class parameters, and turned into genome attributes. Only the GFF3 file is a positional argument. Next, the library implements a ‘gene’ class as a nested ‘genome’ class. Nested classes are generally discouraged in Python except in particular cases. In this library, ‘gene’ objects can only exist within a ‘genome’ object, and the nested approach allows the inheritance of genome attributes shared by all genes in a single step with no intervention of the user. This ‘gene’ class is instantiated with the positional argument of an mRNA feature set by the user as representative of the gene (according to the identifiers in the GFF3 file). Optionally, a protein ID linked to the mRNA, or a description of the gene can be included as keyword arguments. If no protein identifier is supplied, protein domains will not be plotted. The next step is the retrieval of genome coordinates of exons and introns from the GFF3 file. The library requires the creation of a GFF3 database with the *gffutils* Python package (https://github.com/daler/gffutils). The function *createGFFdb*() of the library parses a command to create the database, for convenience. The trunk function of the library is *plot*(), which launches the automated analysis for all other internal functions. It collects the necessary information from the ‘genome’ and ‘gene’ attributes. The function takes as arguments the name of the biological sample (optional) for SNP plotting (from the VCF files), and the type of protein domain (optional) to be plotted in the chart (selected from the available types in the InterproScan output file). If no domain type is supplied, Pfam is used by default. Since protein domain data are reported in the InterproScan file relative to the sequence of the protein, protein coordinates are converted into genome coordinates by the *transcriptpos_to_genomepos*() function of the ‘gene’ class. It builds an index of the nucleotide positions of the CDS (from the GFF3 file) circumventing intron areas, another index of the transcript positions of the gene (from the protein sequence), and both indexes are connected. Next, SNP data is recovered from the VCF files by the *getsnppos*() function. It scans all VCF files in the directory path specified by the user, in order to find the file whose sample name matches the sample argument of the function. Next, SNP positions overlapping with the coordinates of the gene object are retrieved with VCFtools [19]. The VCF file must comply with the standard specifications about sample name specification. Once all data have been collected, the library imports *GenomeDiagram* [20] from the *Biopython* package [17] and plots the UTR (if present), exon and intron subfeatures on the bottom track 0, the protein domain topology on track 1, and SNP data on track 2 (Figure 2 as an example). The function tries to detect SNP annotation by SnpEff [21] from the VCF file. If present, SNPs are colored according to their impact.

**Figure 1.**
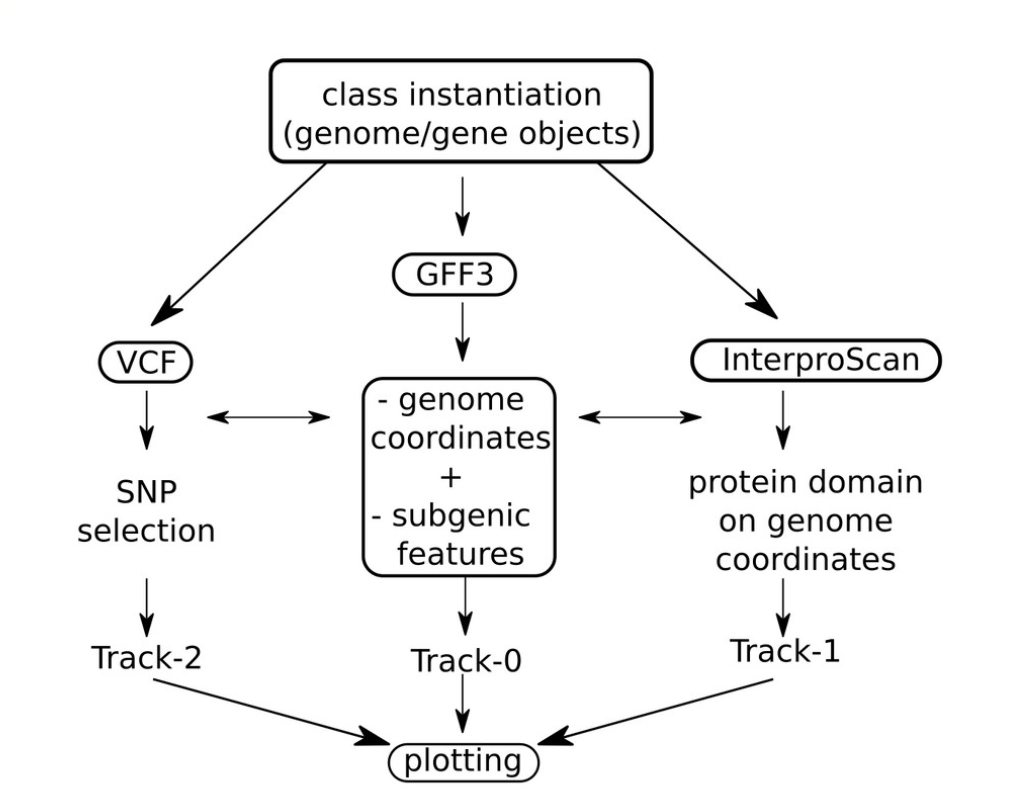
Pipeline workflow

**Figure 2.**
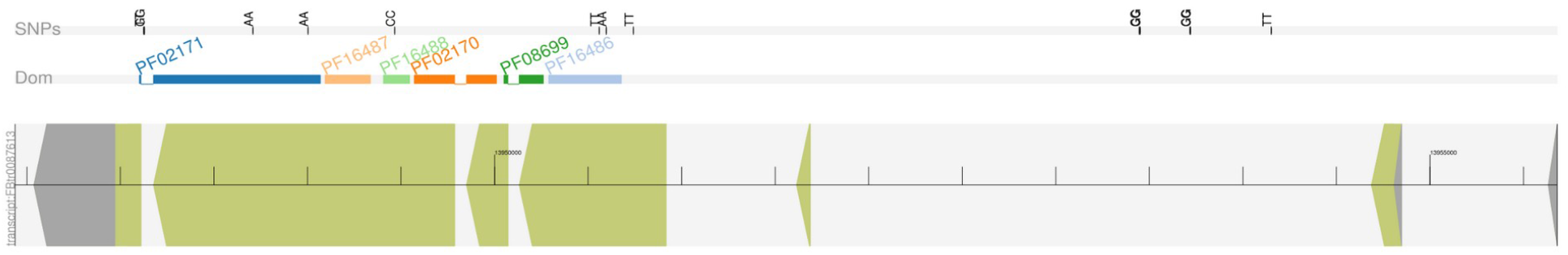
Example of graphical output of the geneplot library. Track 0 (bottom) shows the exon-intron structure (exon=green, UTR=grey). Track 1 shows protein domains. Each domain ID is colored from a color loop sequence. Track 2 (top) shows polymorphism data.

The library is supplied as an open-source package built with *wheel* (https://github.com/pypa/wheel) and published on the Python Package Index (PyPi, https://pypi.org/) repository with the *twine* library (https://github.com/pypa/twine). Package documentation has been created with *Sphinx* (https://www.sphinx-doc.org) and published on the Read The Docs (https://readthedocs.org/) website. A small tutorial has been included in the documentation to set up the library with example data. The code is available from GitHub (https://github.com/) under a GPL v3.0 license, and an automated test for assisting in package development is supplied to be used with the *unittest* Python library (http://docs.python.org/library/unittest.html). Execution and error traceability has been implemented with the *logging* package (https://docs.python.org/3/library/logging.html). The *geneplot* library depends on the external software VCFtools [19], and the non built-in Python libraries *gffutils, Biopython* and *matplotlib* [22].

## Results and discussion

The largest transcripts from 1,351 genes on chromosome 2R from the fruit fly genome (dm6) codifying for a protein larger than 400 amino acids in length were collected, and protein domains were identified with a standalone version of InterproScan (v5.59-91.0). Next, genes were plotted with the *geneplot* library, and genome coordinates of Pfam domains were compared with those reported by the UCSC genome browser for the same genes. In total, 1,147 transcripts with 2,601 recognizable protein domains (2,2 domains/gene on average, 1,101 different Pfam IDs) were analyzed. Figure S1-1 of the supplementary material shows an example of perfect match between both approaches. In several cases, additional Pfam domains were identified or missed in UCSC predictions (e.g. Figure S1-2). This might represent different versions of the Pfam database used by UCSC genome browser and *geneplot*. Since this problem does not relate to the genome coordinate conversion analysis, those cases were discarded and no longer considered for the comparison. Among cases where the same domains were identified, one source of discrepancy consisted in differences that were magnified due to the small length of exons (Figure S1-3 serves as an example). Other times, discrepancy came from the strategy of domain mapping followed by the UCSC tools, which use all available exons of a gene model as a template. Conversely, *geneplot* only uses transcript-associated exons (see Figure S1-4 for another example). Figure S1-5 shows an interesting example of perfect match between domain and exon topology that could potentially promote modularization in new gene biogenesis during evolution under the point of view of the exon shuffling theory. Despite all the aforementioned discrepancies, bidirectional coverage comparison of both data sets agreed in 98.7% and 98.3% (Figure 3), suggesting an equivalent performance. The approach allows taking advantage of raw identifications from InterproScan and the consideration in one single step of the broad range of domain signatures it reports, not just Pfam, but also CDD, Coils, Gene3D, Hamap, MobiDBLite, PANTHER, PIRSF, PRINTS, ProSitePatterns, ProSiteProfiles, SFLD, SMART, SUPERFAMILY and TIGRFAM. Protein signatures corresponding to extremely short domains or even only functional residues (such as those involved in catalysis, folding or protein-ligand interaction) are of special relevance with regard to the applicability of coordinate conversion, since mapping approaches are probably less suitable to correctly transform those protein positions into genomic ranges, due to the small signature length.

**Figure 3.**
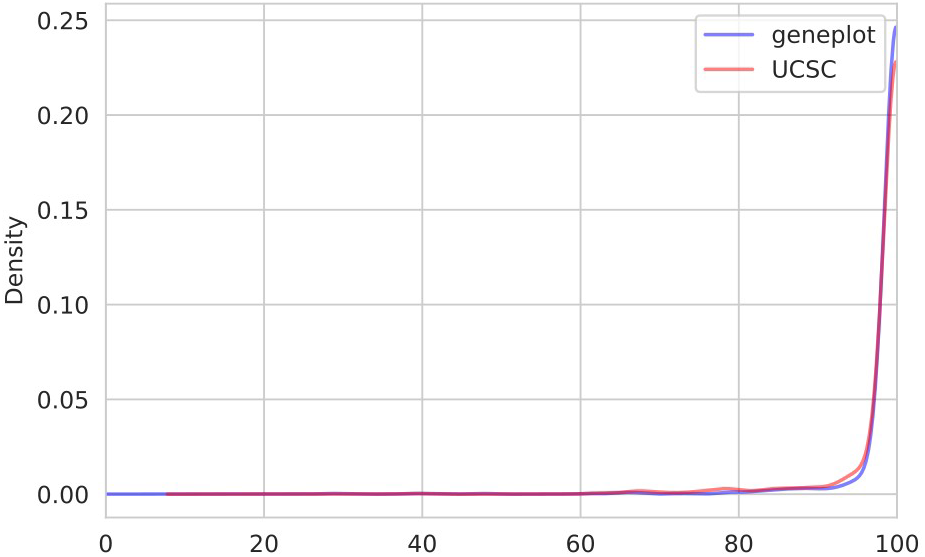
Density plot of protein domain coverages. Genome areas corresponding to protein domains identified by either geneplot (violet) or UCSC (pink) were compared with those that were found by the other method in a set of 1,147 genes. Coverage was expressed as percentage of nucleotides identified as a protein domain by both methods.

## Supplementary material

**Figure S1**

## Bibliography

[1] Y. Wang, H. Zhang, H. Zhong, and Z. Xue, “Protein domain identification methods and online resources,” Comput. Struct. Biotechnol. J., vol. 19, pp. 1145–1153, Jan. 2021, doi: 10.1016/j.csbj.2021.01.041.

[2] Y. Márquez et al., “ExOrthist: a tool to infer exon orthologies at any evolutionary distance,” Genome Biol., vol. 22, no. 1, p. 239, Aug. 2021, doi: 10.1186/s13059-021-02441-9.

[3] M. Liu and A. Grigoriev, “Protein domains correlate strongly with exons in multiple eukaryotic genomes – evidence of exon shuffling?,” Trends Genet., vol. 20, no. 9, pp. 399–403, Sep. 2004, doi: 10.1016/j.tig.2004.06.013.

[4] C. P. Ponting and R. R. Russell, “The Natural History of Protein Domains,” Annu. Rev. Biophys. Biomol. Struct., vol. 31, no. 1, pp. 45–71, Jun. 2002, doi: 10.1146/annurev.biophys.31.082901.134314.

[5] B. Contreras-Moreira, P. F. Jonsson, and P. A. Bates, “Structural Context of Exons in Protein Domains: Implications for Protein Modelling and Design,” J. Mol. Biol., vol. 333, no. 5, pp. 1045–1059, Nov. 2003, doi: 10.1016/j.jmb.2003.09.023.

[6] S. Yousefi et al., “A SNP panel for identification of DNA and RNA specimens,” BMC Genomics, vol. 19, no. 1, p. 90, Jan. 2018, doi: 10.1186/s12864-018-4482-7.

[7] M. W. Ganal, T. Altmann, and M. S. Röder, “SNP identification in crop plants,” Curr. Opin. Plant Biol., vol. 12, no. 2, pp. 211–217, Apr. 2009, doi: 10.1016/j.pbi.2008.12.009.

[8] H. K. Koopaee and A. E. Koshkoiyeh, “SNPs genotyping technologies and their applications in farm animals breeding programs: review,” Braz. Arch. Biol. Technol., vol. 57, pp. 87–95, Feb. 2014, doi: 10.1590/S1516-89132014000100013.

[9] K. K. Amoako, M. C. Thomas, T. W. Janzen, and N. Goji, “Rapid SNP Detection and Genotyping of Bacterial Pathogens by Pyrosequencing,” Methods Mol. Biol. Clifton NJ, vol. 1492, pp. 203–220, 2017, doi: 10.1007/978-1-4939-6442-0_15.

[10] A. Han, H. J. Kang, Y. Cho, S. Lee, Y. J. Kim, and S. Gong, “SNP@Domain: a web resource of single nucleotide polymorphisms (SNPs) within protein domain structures and sequences,” Nucleic Acids Res., vol. 34, no. Web Server issue, pp. W642–W644, Jul. 2006, doi: 10.1093/nar/gkl323.

[11] P. C. Ng and S. Henikoff, “SIFT: Predicting amino acid changes that affect protein function,” Nucleic Acids Res., vol. 31, no. 13, pp. 3812–3814, Jul. 2003, doi: 10.1093/nar/gkg509.

[12] D. N. Cooper, “Functional intronic polymorphisms: Buried treasure awaiting discovery within our genes,” Hum. Genomics, vol. 4, no. 5, pp. 284–288, Jun. 2010, doi: 10.1186/1479-7364-4-5-284.

[13] W. J. Kent et al., “The Human Genome Browser at UCSC,” Genome Res., vol. 12, no. 6, pp. 996–1006, Jun. 2002, doi: 10.1101/gr.229102.

[14] J. Rainer, L. Gatto, and C. X. Weichenberger, “ensembldb: an R package to create and use Ensembl-based annotation resources,” Bioinformatics, vol. 35, no. 17, pp. 3151–3153, Sep. 2019, doi: 10.1093/bioinformatics/btz031.

[15] M. Lawrence et al., “Software for Computing and Annotating Genomic Ranges,” PLOS Comput. Biol., vol. 9, no. 8, p. e1003118, Aug. 2013, doi: 10.1371/journal.pcbi.1003118.

[16] J. E. Stajich et al., “The Bioperl toolkit: Perl modules for the life sciences,” Genome Res., vol. 12, no. 10, pp. 1611–1618, Oct. 2002, doi: 10.1101/gr.361602.

[17] P. J. A. Cock et al., “Biopython: freely available Python tools for computational molecular biology and bioinformatics,” Bioinformatics, vol. 25, no. 11, pp. 1422–1423, Jun. 2009, doi: 10.1093/bioinformatics/btp163.

[18] P. Jones et al., “InterProScan 5: genome-scale protein function classification,” Bioinformatics, vol. 30, no. 9, pp. 1236–1240, May 2014, doi: 10.1093/bioinformatics/btu031.

[19] P. Danecek et al., “The variant call format and VCFtools,” Bioinformatics, vol. 27, no. 15, pp. 2156–2158, Aug. 2011, doi: 10.1093/bioinformatics/btr330.

[20] L. Pritchard, J. A. White, P. R. J. Birch, and I. K. Toth, “GenomeDiagram: a python package for the visualization of large-scale genomic data,” Bioinforma. Oxf. Engl., vol. 22, no. 5, pp. 616–617, Mar. 2006, doi: 10.1093/bioinformatics/btk021.

[21] P. Cingolani et al., “A program for annotating and predicting the effects of single nucleotide polymorphisms, SnpEff,” Fly (Austin), vol. 6, no. 2, pp. 80–92, Apr. 2012, doi: 10.4161/fly.19695.

[22] J. D. Hunter, “Matplotlib: A 2D Graphics Environment,” Comput. Sci. Eng., vol. 9, no. 03, pp. 90–95, May 2007, doi: 10.1109/MCSE.2007.55.

