## Supplemental Figure 1 for "Geneplot: a coordinate conversion approach for graphical representation of protein domain data on the exon-intron structure of a gene"

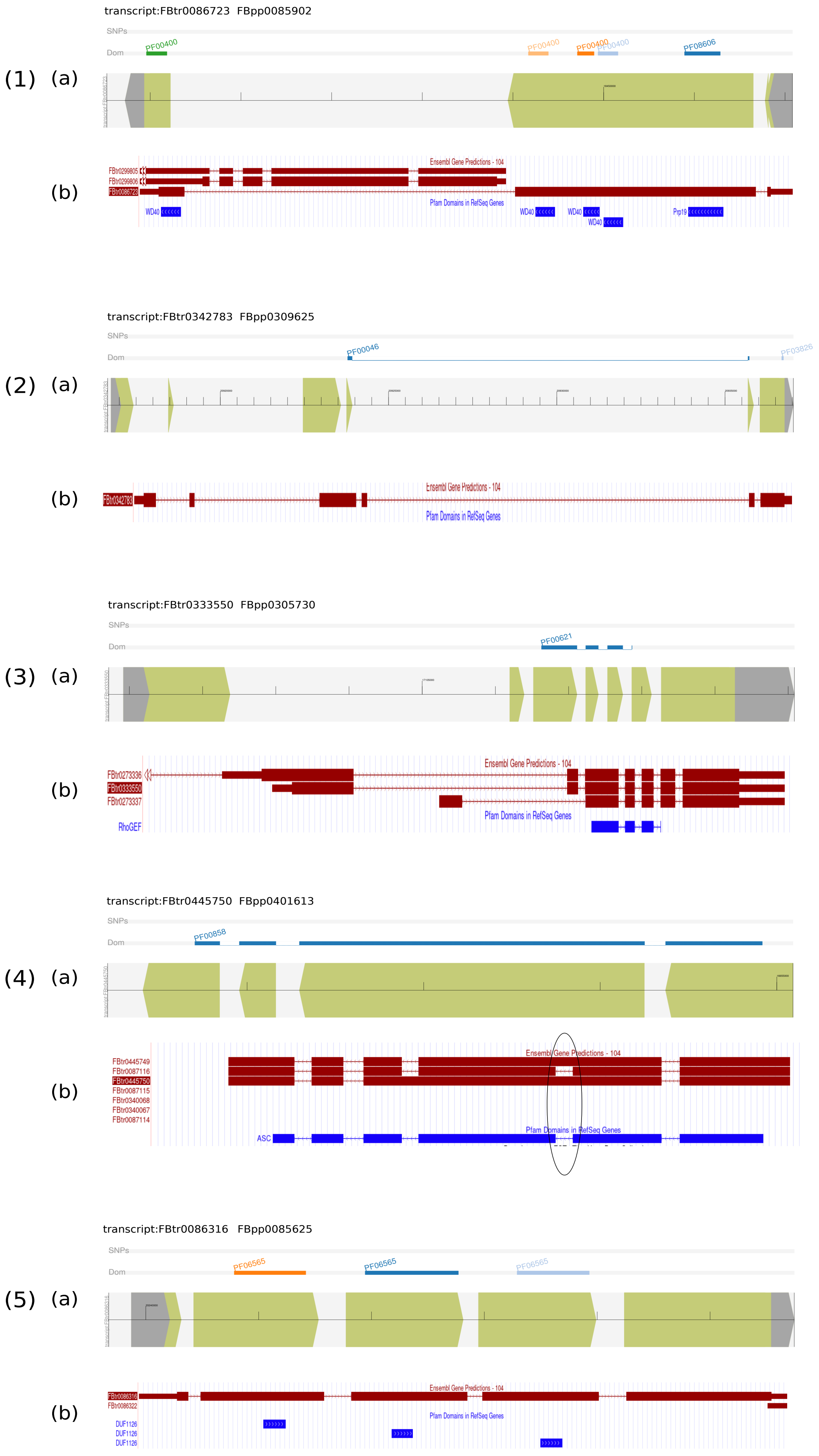


Figure S1. Comparison of genomic coordinates calculation by the UCSC genome browser and the *geneplot* library to represent protein domains over the genome sequence. For each example, a plot output from *geneplot* (a) and a screenshot from the UCSC genome browser (b) are depicted. (1) Perfect match of genome coordinates calculated by both approaches. (2) Discrepancy in Pfam domain identification on raw protein sequences (before coordinate calculation) potentially due to different versions of Pfam used by both approaches. These types of cases were discarded for the comparison. (3) Discrepancy in 3 nucleotides in one exon of 5 nucleotides in length, resulting in a 60% difference. This example explains how in some cases percentage differences were magnified. (4) Discrepancy in the number of exons spanning the protein domain between UCSC and *geneplot* results. UCSC uses as template all possible exons available for a single gene model. Conversely, geneplot only uses transcript-associated exons. (5) The same domains were identified by both approaches, but with different domain length on raw protein sequences (before coordinate calculation). This result is probably due to different versions of Pfam used by both approaches.
